# Topology-aware thermodynamics for DNA probe design under fixed stringency Retained paired boxes link mismatch placement, nearest-neighbor stability and room-temperature diagnostic specificity

**DOI:** 10.64898/2026.06.10.731169

**Authors:** Ivan Brukner, Maja Krajinovic

## Abstract

DNA probe specificity is usually screened by mismatch number, melting temperature and full-duplex nearest-neighbor free energy. These scalar quantities are necessary but obscure the spatial organization of pairing: an off-target can retain one long paired box or split the same matches into short fragments. We define thermodynamics without topology as scalar full-duplex nearest-neighbor DeltaG and thermodynamics with topology as redistribution of nearest-neighbor contributions over retained paired boxes. The practical score family begins with edge-corrected effective complementary islands, extends to the count Nbox(L,k) of contiguous boxes of length at least k, and culminates in Zbox,NN(k,T), a retained-box partition weighted by nearest-neighbor stability. Reanalysis of published mismatch-probe datasets shows why the distinction matters: distributed or staggered mismatches are far more disruptive than clustered mismatches at comparable burden; 50-mer mismatch probes show distribution- and temperature-dependent residual signal; maximum perfect-match length explains a substantial fraction of 60-mer mismatch signal ratios; and zipper/partition-function modeling explains position-dependent mismatch discrimination. An Affymetrix fixed-mismatch subset and an ambient-temperature HPV probe panel provide independent control and diagnostic context. The framework offers an auditable rule for fixed-stringency probe design: preserve intended retained continuity, fragment the strongest off-target retained box, then validate in the final buffer and readout.

## INTRODUCTION

Probe design traditionally relies on GC content, melting temperature, mismatch count and full-duplex nearest-neighbor DeltaG. These quantities are essential, but they become incomplete when one probe must discriminate closely related amplicons under one fixed hybridization condition. Two off-target alignments can contain the same number of mismatches but very different retained continuity: one may preserve a long uninterrupted paired box, while another may split the same matched bases into short fragments.

This problem is acute in room-temperature and point-of-care nucleic-acid assays. PCR can create abundant amplicons, lateral-flow strips offer limited opportunity for temperature tuning or stringent washing, and weak off-target hybridization can still generate a visible band. Under these constraints, a useful probe is not simply the strongest exact complement. It is a sequence that preserves intended retained pairing while fragmenting retained paired boxes in the closest non-intended targets.

The diagnostic basis for this problem was established by earlier probe-selection and ambient-temperature HPV typing studies [9-11]. Those studies showed that closely related viral sequences can be resolved under shared, non-denaturing, PCR-compatible conditions. The present work does not claim that those studies used ECI or Zbox,NN explicitly; rather, it uses them as the applied fixed-stringency context for a general topology-aware thermodynamic framework.

The central distinction is not topology versus thermodynamics. It is thermodynamics without topology versus thermodynamics with topology. Thermodynamics without topology collapses an aligned duplex into a scalar full-duplex DeltaG. Thermodynamics with topology keeps the linear arrangement of retained pairing and applies nearest-neighbor logic to retained contiguous boxes that can nucleate, persist or rehybridize under a given stringency.

**Figure 1.**
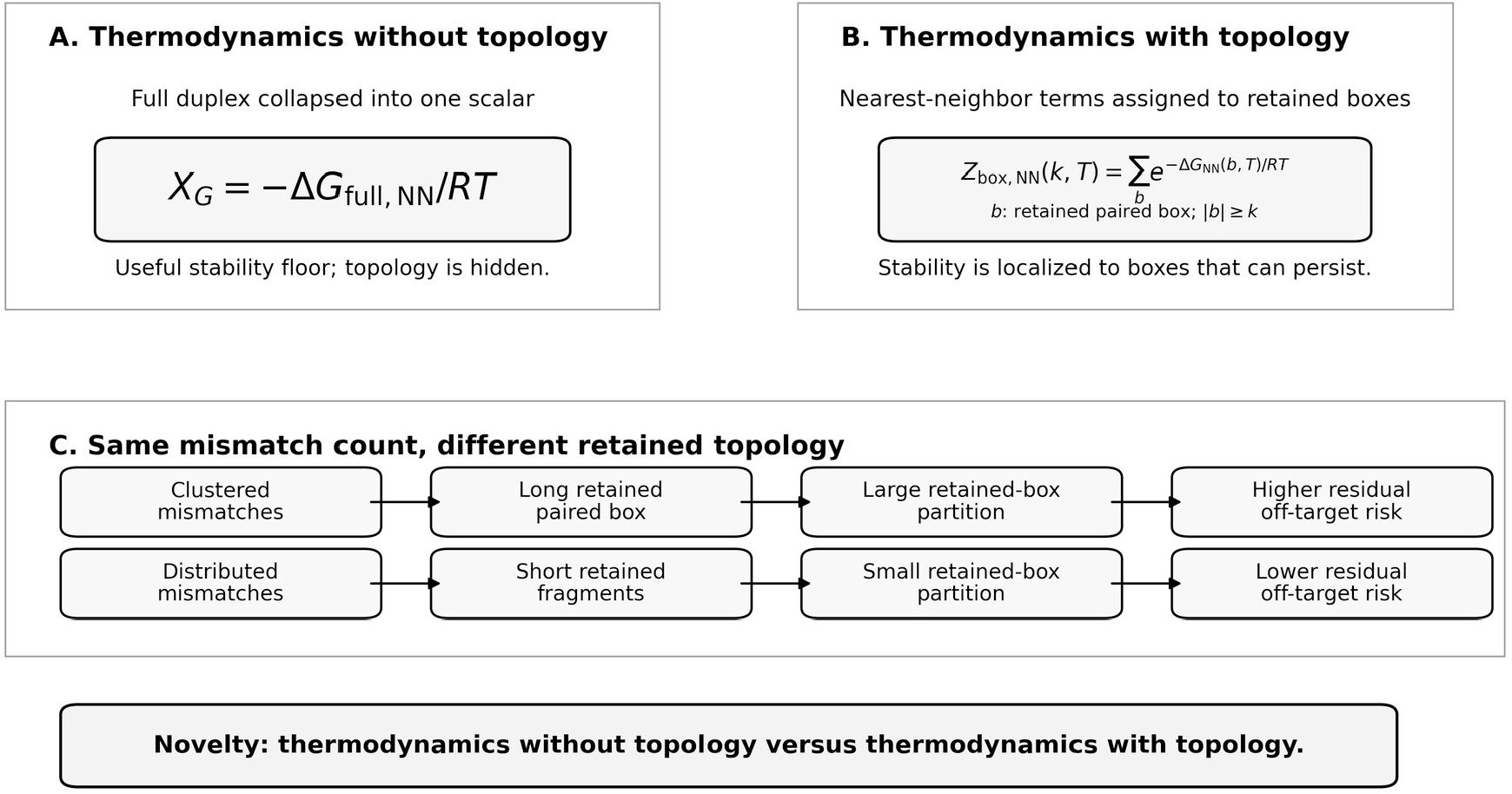
Topology-aware thermodynamics for probe specificity. Thermodynamics without topology versus thermodynamics with topology. Scalar full-duplex DeltaG gives a global stability value, whereas topology-aware thermodynamics reports whether stable contributions remain as one long retained paired box or are fragmented into shorter boxes.

## MATERIALS AND METHODS

### Definitions and edge-corrected retained islands

A formal matched island is an uninterrupted run of Watson-Crick matches in a probe-target alignment. A retained island is the formal island after subtracting one base for each mismatch- or gap-exposed edge. For island i with formal length Li and exposed-edge count ei, the retained core is ci = max(0, Li - ei). Retained cores below an assay-specific threshold are ignored for design acceptance.

The practical effective complementary island score is S_ECI = sum_i ECI_i^2. The square is a convex ranking rule, not a new free-energy law. It distinguishes one long retained island from several short fragments with the same total number of matches.

The combinatorial bridge to the square rule is Nbox(L,k) = (L-k+1)(L-k+2)/2 for L >= k, and zero otherwise. This counts the number of contiguous retained paired boxes of length at least k within one retained island of length L. When L is larger than k, Nbox grows approximately quadratically with L. The thermodynamic retained-box partition is Zbox,NN(k,T) = sum_b exp[-DeltaG_NN(b,T)/RT] over retained boxes b with length at least k.

### Datasets and evidence layers

The manuscript separates evidence roles. Same-burden external microarray studies provide the strongest tests of topology because mismatch number or gross thermodynamic burden is held similar while placement changes. The Seringhaus/Gerstein tiling-array study supplies a centered-versus-staggered mismatch contrast; the Deng/Zhou 50-mer PM/MM study supplies even-versus-random mismatch placement and temperature thresholds; the Rennie/Hoyle long-oligonucleotide CGH study supplies maximum perfect-match length as a retained-continuity predictor; and the Naiser study supplies the double-ended zipper/partition-function mechanism. Hooyberghs/Carlon and Hadiwikarta-type microarray thermodynamic work provide the scalar nearest-neighbor baseline and its limits.

The Affymetrix fixed-mismatch subset is used as a small independent control in which topology is compared at fixed mismatch count. The ambient-temperature HPV probe panel is used as an applied fixed-stringency diagnostic audit, not as the primary statistical proof, because the original panel consisted of empirically functional probes. All reconstructed tables are supplied as Supplementary Tables S1-S11 and CSV files.

### Statistical analyses and model comparisons

Where row-level data were available, correlations were computed between signal endpoints and retained-continuity descriptors such as Lmax, S_ECI and Nbox. Where only published aggregate values were available, the values were reconstructed from reported equations or figures and labelled as literature-reported aggregates. The main text emphasizes same-burden contrasts because pooled regressions can make topology appear merely incremental when mismatch count, GC content and scalar DeltaG already capture much of the easy variance.

The central model contrast is DeltaG_full,NN alone versus DeltaG_full,NN plus retained-box topology, or the direct retained-box partition Zbox,NN when sequence-level nearest-neighbor box weights are available. All claims about retrospective analyses are treated as hypothesis-supporting and design-auditing rather than prospective clinical validation.

### Probe-design algorithm for fixed-stringency assays

1. Define intended amplicons and a near-neighbor off-target panel.
2. Generate exact-complement, shifted, shortened and limited intentional-mismatch probe candidates.
3. Apply conventional filters: GC content, Tm, full-duplex DeltaG, self-structure and synthesis constraints.
4. Align candidates against intended and non-intended windows and compute retained islands, S_ECI, Nbox and, where possible, Zbox,NN.
5. Rank off-targets by retained-box risk, not mismatch count alone.
6. Redesign locally to split the highest-risk off-target retained box while preserving intended retained continuity.
7. Validate under the final buffer, temperature, surface chemistry, flow time and readout threshold.

## RESULTS

### Same-burden controls reveal large topology effects

The most persuasive evidence comes from datasets in which mismatch burden is similar but mismatch placement differs. In the Seringhaus/Gerstein 36-mer tiling-array study, centered mismatches were much less disruptive than staggered mismatches: the fitted slopes correspond to approximately 7% loss of perfect-match intensity per centered mismatch and approximately 21% per staggered mismatch. Thus, distributing mismatches along the oligonucleotide is about three-fold more disruptive than adding adjacent mispairs under the reported conditions [3].

The Deng/Zhou 50-mer PM/MM study provides an independent topology and stringency control. At the same mismatch count, evenly distributed mismatches reduced residual signal more than randomly distributed mismatches. For four mismatches, the reported average relative signal was approximately 0.11 for evenly distributed mismatches and 0.26 for randomly distributed mismatches. The same study also showed a temperature-dependent threshold: three, four and five evenly distributed mismatches were needed at 50, 45 and 42 C, respectively, to suppress residual signal to their practical low-signal criterion [4].

The Rennie/Hoyle 60-mer CGH study shows the retained-length principle in a different system: maximum perfect-match length explained about 43% of the variance in log2 signal ratios between probes with one and two mismatches, and mismatch position contributed more variance than substitution type [5]. Naiser et al. provide the mechanistic explanation: partial duplex states in a double-ended zipper/partition-function model can reproduce position-dependent mismatch discrimination [6].

**Figure 2.**
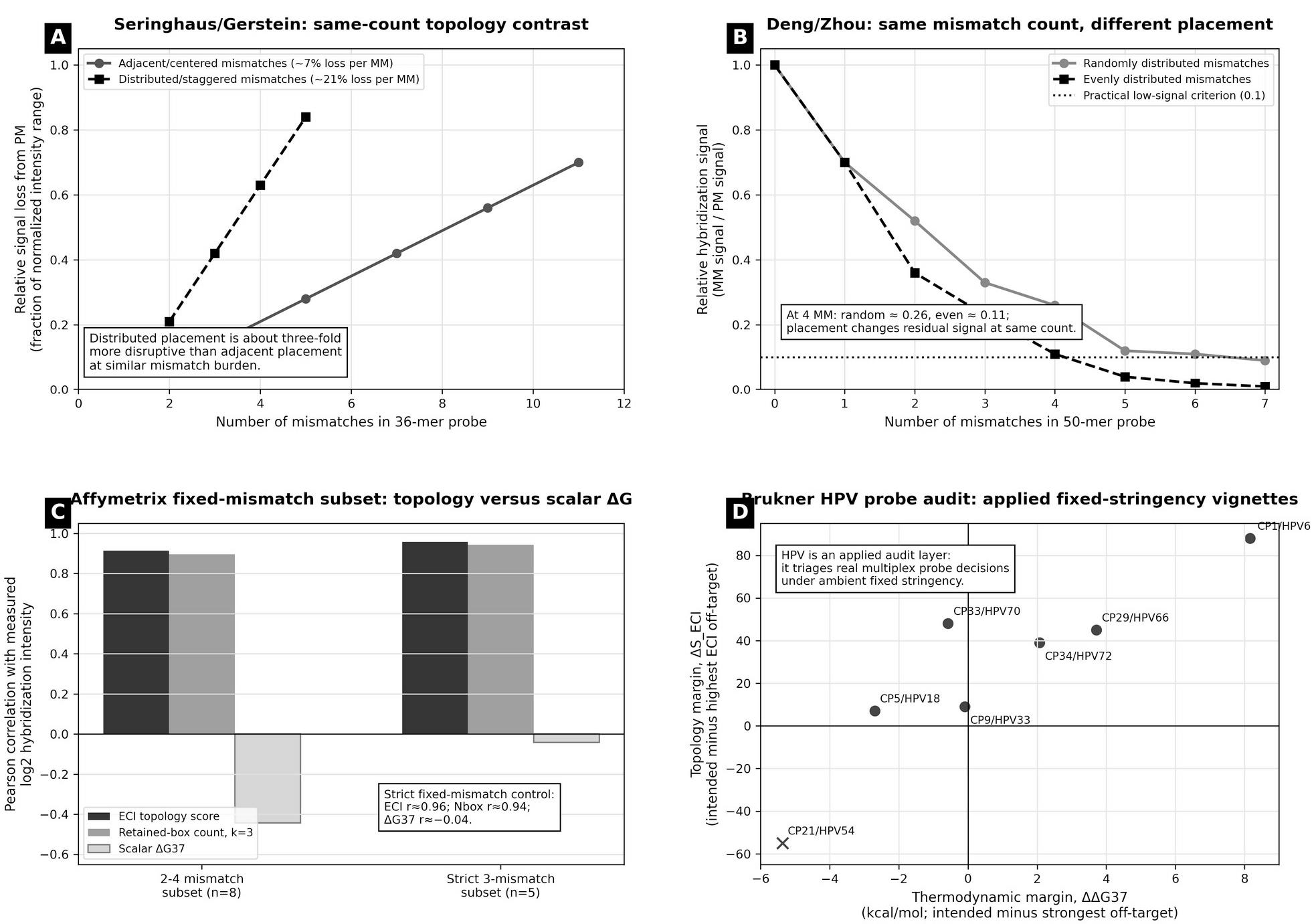
Extreme controls plus retained Affymetrix and HPV evidence. Same-burden topology evidence plus fixed-mismatch and ambient diagnostic context. Full numerical values are provided in Supplementary Tables S2-S10.

### From square-like retained-island scoring to topology-aware thermodynamics

The retained-island score S_ECI is useful because it is visible and actionable, but its square term can be explained more directly by Nbox. A retained island of length L contains a growing number of sub-boxes of length at least k. Once L exceeds k, the number of possible retained boxes grows approximately quadratically. This makes one long off-target island much more risky than the same number of matched bases split into fragments.

The thermodynamic extension, Zbox,NN, replaces unweighted box counting with nearest-neighbor weights for each retained box. This preserves the nearest-neighbor framework while exposing where the stabilizing terms are located along the duplex. In this sense, topology-aware thermodynamics is not an alternative to nearest-neighbor thermodynamics; it is a redistribution of nearest-neighbor contributions over the retained paired boxes that matter under a specified stringency.

### Fixed-mismatch and ambient diagnostic application layers

The Affymetrix fixed-mismatch subset is retained as a small but clean control. In the 2-4 mismatch subset, S_ECI and Nbox(k=3) correlated with log2 intensity at r = 0.915 and r = 0.896, respectively. In the strict 3-mismatch subset, S_ECI and Nbox(k=3) remained strongly associated with intensity, while DeltaG37 alone was weak or negative (Supplementary Tables S8-S9).

The ambient-temperature HPV probe-selection and typing studies supply the applied diagnostic anchor. The 39-probe HPV panel should not be used as the main statistical proof because those probes were empirically functional, but it is highly useful as a fixed-stringency audit. It shows how scalar DeltaG and retained topology can nominate different risk drivers and guide local probe redesign under ambient, PCR-compatible conditions (Supplementary Table S10).

### Stringency calibrates the effective box-length threshold

The minimum retained-box length k_eff is not universal. It changes with temperature, salt and formamide concentration, probe chemistry, target concentration, surface chemistry, flow time and readout threshold. Deng/Zhou provide a direct calibration example in 50% formamide: the practical mismatch threshold increased from three mismatches at 50 C to five mismatches at 42 C. This is consistent with the interpretation that lower stringency allows shorter residual off-target boxes to contribute to signal.

Room-temperature post-PCR lateral-flow assays represent a particularly conservative setting: amplicon concentration can be high, hybridization is brief, and wash stringency is limited. Therefore, probe candidates should be screened for long retained off-target boxes at a low-stringency k_eff, then validated empirically in the final strip and buffer.

**Figure 3.**
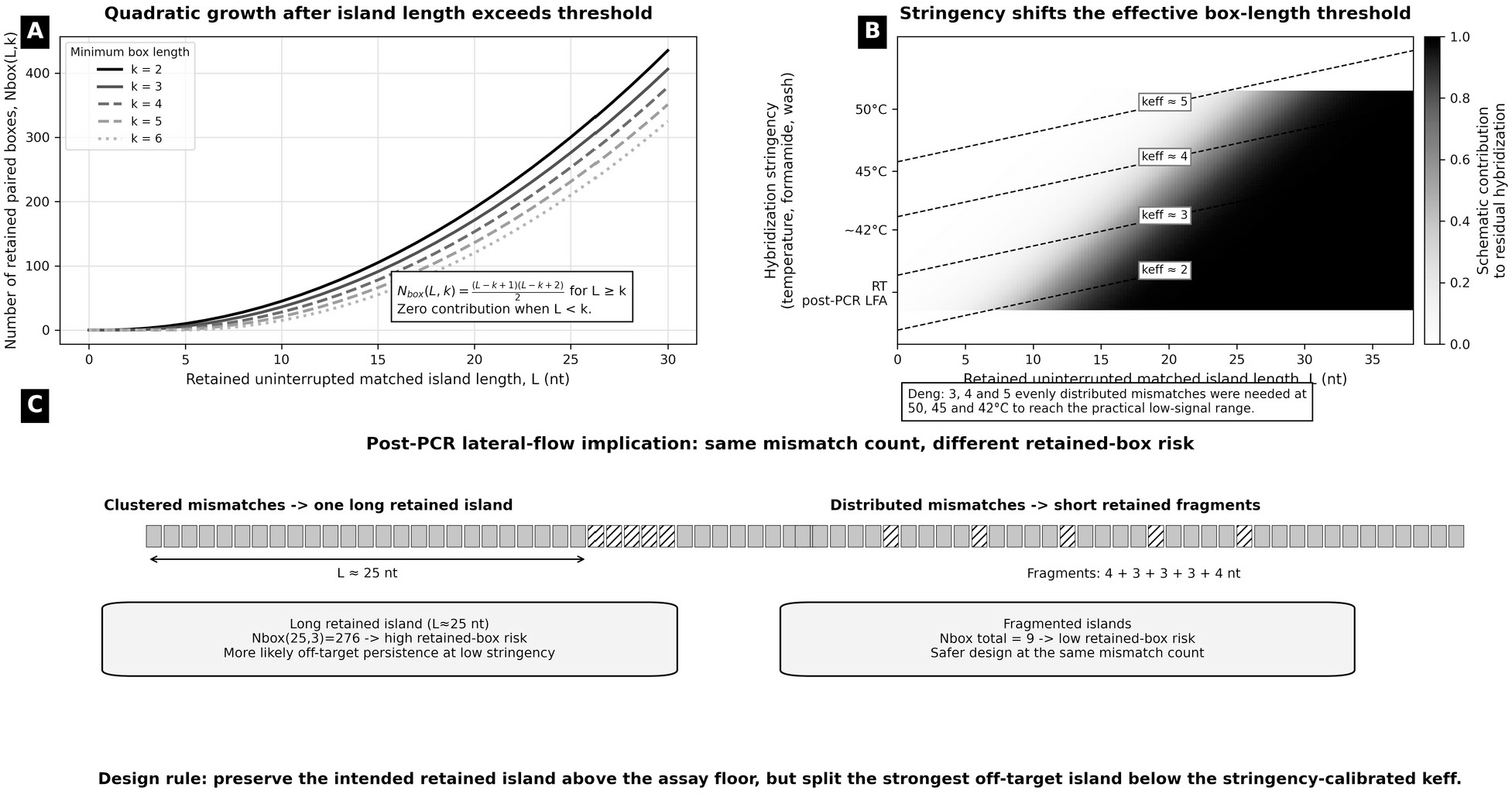
Retained-island length, effective box threshold and room-temperature stringency. Retained-island length, effective box threshold and hybridization stringency. Nbox(L,k) provides the combinatorial basis for square-like retained-island behavior, and k_eff is calibrated by assay stringency rather than fixed universally.

### Design workflow for room-temperature post-PCR lateral-flow probes

The design objective is not maximum on-target affinity alone. The objective is to preserve the intended retained island above the assay signal floor while reducing the strongest non-intended retained island below the stringency-calibrated threshold. This produces a local design action: shift the probe, shorten an end, or introduce a limited intentional mismatch to fragment the highest-risk off-target box.

The same logic naturally extends to broad-range 16S bacterial identification, where conserved primers amplify a mixed amplicon family and species-specific information lies between primer sites. This manuscript does not claim completed 16S validation. It identifies 16S post-PCR lateral-flow identification as a direct next benchmark for the same topology-aware design rule.

**Figure 4.**
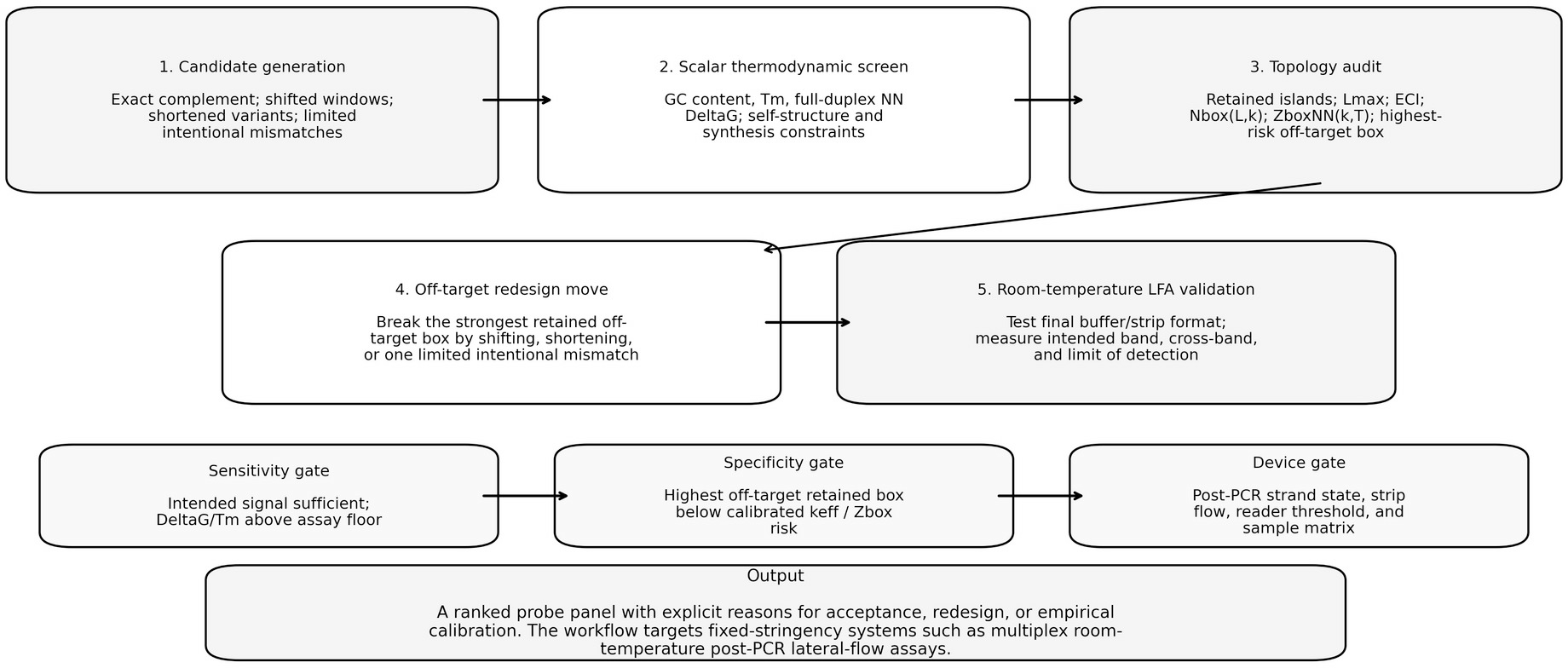
Topology-aware probe-design logic for post-PCR lateral-flow hybridization. Probe-design logic for topology-aware fixed-stringency hybridization. Scalar thermodynamics is retained as a sensitivity gate, while retained-box topology functions as a specificity gate that identifies exactly which off-target island should be disrupted.

## DISCUSSION

The data support a specific claim: topology becomes decisive when conventional variables are ambiguous. In pooled datasets, scalar DeltaG, mismatch count and GC content can explain much of the signal. In same-count or position-controlled settings, however, topology reveals why one alignment survives and another fails: retained paired boxes are either preserved or fragmented.

This framework also explains why the square-like ECI score is useful without claiming that free energy itself is quadratic. The square-like behavior arises because the number of possible retained boxes grows approximately quadratically once a retained island exceeds the effective threshold. Zbox,NN then supplies the thermodynamic extension by weighting each retained box by nearest-neighbor stability.

The method should be interpreted as a design and audit layer, not a universal physical law. It does not replace full thermodynamics, BLAST or alignment screening, self-structure analysis, target accessibility assessment, surface/matrix effects or empirical testing. Instead, it provides a missing, interpretable variable: where the off-target still retains a dangerous paired box.

For fixed-stringency room-temperature assays, especially post-PCR lateral-flow formats, this variable is practically important. High amplicon load and limited washing can allow weak off-target boxes to produce visible lines. Topology-aware thermodynamics therefore supplies a rational presynthesis filter and a concrete redesign instruction.

## DATA AVAILABILITY

All reconstructed aggregate tables, Affymetrix fixed-mismatch calculations, HPV audit rows, plotted values, figure files, CSV files and the Excel workbook Supplementary_Numerical_Tables_S1_S11.xlsx are included as Supplementary Data with this submission. External raw data remain with the cited publications and public repositories described in those references. Before final submission, the Supplementary Data directory should be deposited in a permanent repository such as Zenodo or Figshare and the DOI inserted here. No new biological specimens or novel sequences were generated.

## SUPPLEMENTARY DATA

Supplementary Data are available at NAR Online. The Supplementary Information contains scoring definitions, derivations, evidence-layer notes, numerical tables S1-S11, source-role audit, stringency calibration and data inventory. Machine-readable CSV copies and a single Excel workbook containing all numerical tables are included in the Supplementary Data directory.

## ACKNOWLEDGEMENTS

The authors acknowledge the scientific foundation provided by earlier probe-selection and HPV typing work, and the public PM/MM microarray studies that made independent topology comparisons possible. Any AI-assisted drafting or coding support should be disclosed according to journal policy in the cover letter and acknowledgements if used for the final submitted version.

## AUTHOR CONTRIBUTIONS

Conceptualization: I.B. and M.K.; methodology: I.B.; formal analysis: I.B.; investigation: I.B.; data curation: I.B.; visualization: I.B.; writing - original draft: I.B.; writing - review and editing: I.B. and M.K.; supervision: M.K. All authors have read and approved the manuscript draft.

## REFERENCES

1. SantaLucia J Jr, Hicks D. The thermodynamics of DNA structural motifs. Annu Rev Biophys Biomol Struct. 2004;33:415–440.

2. Yakovchuk P, Protozanova E, Frank-Kamenetskii MD. Base-stacking and base-pairing contributions into thermal stability of the DNA double helix. Nucleic Acids Res. 2006;34:564–574.

3. Seringhaus M, Rozowsky J, Royce T, Nagalakshmi U, Jee J, Snyder M, Gerstein M. Mismatch oligonucleotides in human and yeast: guidelines for probe design on tiling microarrays. BMC Genomics. 2008;9:635.

4. Deng Y, He Z, Van Nostrand JD, Zhou J. Design and analysis of mismatch probes for long oligonucleotide microarrays. BMC Genomics. 2008;9:491.

5. Rennie C, Noyes HA, Kemp SJ, Hulme H, Brass A, Hoyle D. Strong position-dependent effects of sequence mismatches on signal ratios measured using long oligonucleotide microarrays. BMC Genomics. 2008;9:317.

6. Naiser T, Kayser J, Mai T, Michel W, Ott A. Position dependent mismatch discrimination on DNA microarrays - experiments and model. BMC Bioinformatics. 2008;9:509.

7. Hooyberghs J, Van Hummelen P, Carlon E. The effects of mismatches on hybridization in DNA microarrays: determination of nearest neighbor parameters. arXiv:1001.0653.

8. Hadiwikarta WW, Walter JC, Hooyberghs J, Carlon E. Probing hybridization parameters from microarray experiments: nearest-neighbor model and beyond. Nucleic Acids Res. 2012;40:e138.

9. Harbig J, Sprinkle R, Enkemann SA. A sequence-based identification of the genes detected by probesets on the Affymetrix U133 plus 2.0 array. Nucleic Acids Res. 2005;33:e31.

10. Wang Y, Miao ZH, Pommier Y, Kawasaki ES, Player A. Characterization of mismatch and high-signal intensity probes associated with Affymetrix GeneChips. Bioinformatics. 2007;23:2088–2095.

11. Brukner I, El-Ramahi R, Sawicki J, Gorska-Flipot I, Krajinovic M, Labuda D. Hybridization assay performed at ambient temperature for typing high-risk human papillomaviruses. J Clin Virol. 2007;39:113–118.

12. Brukner I, El-Ramahi R, Gorska-Flipot I, Krajinovic M, Labuda D. An in vitro selection scheme for oligonucleotide probes to discriminate between closely related DNA sequences. Nucleic Acids Res. 2007;35:e66.

13. Brukner I, Krajinovic M, Dascal A, Labuda D. A protocol for the in vitro selection of specific oligonucleotide probes for high-resolution DNA typing. Nat Protoc. 2007;2:2806–2819.

14. Demidov VV, Frank-Kamenetskii MD. Two sides of the coin: affinity and specificity of nucleic acid interactions. Trends Biochem Sci. 2004;29:62–71.

15. Letowski J, Brousseau R, Masson L. Designing better probes: effect of probe size, mismatch position and number on hybridization in DNA oligonucleotide microarrays. J Microbiol Methods. 2004;57:269–278.

16. Owczarzy R, Moreira BG, You Y, Behlke MA, Walder JA. Predicting stability of DNA duplexes in solutions containing magnesium and monovalent cations. Biochemistry. 2008;47:5336–5353.

17. Hertel S, Spinney RE, Xu SY, Ouldridge TE, Morris RG, Lee LK. The stability and number of nucleating interactions determine DNA hybridization rates in the absence of secondary structure. Nucleic Acids Res. 2022;50:7829–7841.

18. Baker GC, Smith JJ, Cowan DA. Review and re-analysis of domain-specific 16S primers. J Microbiol Methods. 2003;55:541–555.

19. Klindworth A, Pruesse E, Schweer T, Peplies J, Quast C, Horn M, Glockner FO. Evaluation of general 16S ribosomal RNA gene PCR primers for classical and next-generation sequencing-based diversity studies. Nucleic Acids Res. 2013;41:e1.

