## Supplement material for "Topology-aware thermodynamics for DNA probe design under fixed stringency Retained paired boxes link mismatch placement, nearest-neighbor stability and room-temperature diagnostic specificity"

**Author:** Ivan Brukner

**Contents:** Supplementary Information and Numerical Tables S1-S11.

**Repository link:** https://doi.org/10.5281/zenodo.20613900

This single PDF is prepared for submission as supplementary material with the bioRxiv preprint. Machine-readable CSV/XLSX tables and high-resolution figure files are deposited separately in Zenodo.

### Supplementary Information and Numerical Tables

Topology-aware thermodynamics for DNA probe design under fixed stringency

This file is intended for NAR peer review. It embeds the numerical tables S1-S11 and is accompanied by a machine-readable Excel workbook and CSV files. External raw data remain with the cited publications or public repositories. Reconstructed tables are explicitly labelled as reported aggregate, reconstructed aggregate, row-level illustrative control, or applied audit.

#### Supplementary Note S0. Data provenance

| Supplementary table | Data type | Provenance / use |
|---|---|---|
| Table S1. Definitions and scoring conventions | Embedded table + CSV + Excel worksheet | Author-defined terms used in the manuscript; formulas are defined analytically. |
| Table S2. Evidence-layer inventory | Embedded table + CSV + Excel worksheet | Synthesis table mapping published datasets and internal audit layers to evidence roles. |
| Table S3. Seringhaus centered/staggered mismatch reconstruction | Embedded table + CSV + Excel worksheet | Reconstructed from reported design positions and fitted equations in Seringhaus et al.; topology metrics calculated from the reported mismatch positions. |
| Table S4. Deng even-versus-random mismatch aggregate values | Embedded table + CSV + Excel worksheet | Reported aggregate relative signal values extracted from Deng et al. Figure 1/text; ratio calculated here. |
| Table S5. Deng MPDNN model progression | Embedded table + CSV + Excel worksheet | Reported model correlations from Deng et al.; R2 calculated as R squared. |
| Table S6. Rennie maximum perfect-match-length summary | Embedded table + CSV + Excel worksheet | Published summary values from Rennie et al. with no row-level reanalysis. |
| Table S7. Mechanistic and thermodynamic bridge evidence | Embedded table + CSV + Excel worksheet | Extracted summary evidence from Naiser et al. and Hooyberghs/Carlon; used for interpretation, not a pooled regression. |
| Table S8. Affymetrix fixed-mismatch control rows | Embedded table + CSV + Excel worksheet | Row-level illustrative control retained from the ECI analysis; sequence-resolved source extraction should be deposited for final submission. |
| Table S9. Affymetrix topology and thermodynamic correlation summary | Embedded table + CSV + Excel worksheet | Pearson correlations and p-values computed from Table S8 rows. |
| Table S10. HPV sequence-resolved edge-case audit | Embedded table + CSV + Excel worksheet | Sequence-resolved applied audit from the ambient HPV benchmark. Rows marked DISCREPANCY_CHECK should be verified before relying on them for probe ordering. |
| Table S11. Temperature/stringency calibration table | Embedded table + CSV + Excel worksheet | Calibration logic derived from Deng et al. temperature thresholds and the proposed room-temperature LFA use case. |

#### Supplementary Note S1. Definitions and scoring conventions

Formal matched islands are uninterrupted Watson-Crick matched runs. Retained islands are formal islands corrected by subtracting one base for each mismatch- or gap-exposed edge. The practical ECI score is a design heuristic, Nbox is a combinatorial paired-box count, and Zbox,NN is the thermodynamic retained-box extension.

#### Supplementary Note S2. Nbox derivation

For a retained island of length L, the number of sub-boxes of exact length m is L-m+1. Summing over all m from k through L gives Nbox(L,k) = sum_{m=k}^{L}(L-m+1) = (L-k+1)(L-k+2)/2 for L >= k. For L much larger than k, this grows approximately as one-half of a square, explaining why square-like ECI behavior is useful without claiming quadratic free energy.

### Table S1. Definitions and scoring conventions

Author-defined terms used in the manuscript; formulas are defined analytically.

| Symbol/term | Definition | Use in manuscript |
|---|---|---|
| Formal matched island | Uninterrupted Watson-Crick matched run in an alignment before edge correction | Starting object for topology scoring |
| Retained island | Matched island after subtracting mismatch- or gap-exposed edges | Corrects formal runs for local disruption |
| L | Retained uninterrupted matched island length in nucleotides | Input to Nbox and ECI |
| k or k_eff | Minimum retained paired-box length considered effective under an assay stringency | Calibrated by temperature, buffer, concentration and readout |
| S_ECI | Sum of squared retained island lengths above threshold | Practical visual topology score |
| Nbox(L,k) | Number of contiguous retained paired boxes of length at least k within an island of length L | Combinatorial bridge explaining square-like behavior |
| Zbox,NN(k,T) | Boltzmann-weighted sum over retained boxes using nearest-neighbor DeltaG for each box | Thermodynamics with topology |
| DeltaG_full,NN | Nearest-neighbor free energy of the full aligned duplex | Thermodynamics without topology baseline |

### Table S2. Evidence-layer inventory

Synthesis table mapping published datasets and internal audit layers to evidence roles.

| Evidence layer | Dataset/source | What is controlled or varied | Most relevant result | Role in NAR manuscript |
|---|---|---|---|---|
| External same-count contrast | Seringhaus et al. 36-mer tiling arrays | Adjacent/centered versus staggered/distributed mismatch placement | Centered mismatches: about 7% signal loss per mismatch; staggered: about 21% per mismatch | Large topology effect under similar mismatch burden |
| External PM/MM design | Deng et al. 50-mer probes | Even versus random mismatch placement; temperature 42/45/50 C | Evenly distributed mismatches lower residual signal; practical threshold shifts from 3 to 5 mismatches as temperature decreases | Topology plus stringency calibration |
| Retained-length support | Rennie et al. 60-mer CGH arrays | 1- and 2-mismatch probes paired by maximum perfect-match length | Maximum perfect-match length explained about 43% of log2 signal-ratio variance | Independent retained-island support |
| Mechanistic model | Naiser et al. in-house microarrays | Defect position and sequence context; zipper/partition modeling | Mismatch discrimination mostly determined by defect position and sequence context | Physical bridge to thermodynamics with topology |
| Microarray thermodynamic baseline | Hooyberghs/Carlon and Hadiwikarta et al. | Nearest-neighbor free energy on arrays | Microarray data can be used to infer hybridization thermodynamics; additivity can break down for close mismatches | Thermodynamic foundation and limits |
| Small fixed-mismatch control | Affymetrix fixed-mismatch subset | Same mismatch count with clustered versus distributed topology | Topology metrics remain correlated with signal; DeltaG37 is weak in strict 3-mismatch subset | Independent small control |
| Ambient diagnostic anchor | HPV probe-selection and typing studies | Closely related HPV motifs under shared ambient PCR-compatible conditions | Probe selection and ambient HPV typing establish the diagnostic use case | Room-temperature application anchor |

### Table S3. Seringhaus centered/staggered mismatch reconstruction

Reconstructed from reported design positions and fitted equations in Seringhaus et al.; topology metrics calculated from the reported mismatch positions.

| study | gene | scheme | n_mismatches | mm_positions | fitted_MM_minus_PM | relative_intensity_loss | R2_reported | Lmax | SECI | Nbox_k3 | retained_runs |
|---|---|---|---|---|---|---|---|---|---|---|---|
| Seringhaus | ACT1 | centered | 3 | 17;18;19 | -0.1754 | 0.1754 | 0.995 | 16 | 481 | 196 | 15;16 |
| Seringhaus | ACT1 | centered | 5 | 16;17;18;19;20 | -0.30920000000000003 | 0.30920000000000003 | 0.995 | 15 | 421 | 169 | 14;15 |
| Seringhaus | ACT1 | centered | 7 | 15;16;17;18;19;20;21 | -0.443 | 0.443 | 0.995 | 14 | 365 | 144 | 13;14 |
| Seringhaus | ACT1 | centered | 9 | 14;15;16;17;18;19;20;21;22 | -0.5768 | 0.5768 | 0.995 | 13 | 313 | 121 | 12;13 |
| Seringhaus | ACT1 | centered | 11 | 13;14;15;16;17;18;19;20;21;22;23 | -0.7106 | 0.7106 | 0.995 | 12 | 265 | 100 | 11;12 |
| Seringhaus | ACT1 | staggered | 2 | 12;24 | -0.2168 | 0.2168 | 0.998 | 11 | 302 | 109 | 10;9;11 |
| Seringhaus | ACT1 | staggered | 3 | 12;18;24 | -0.4306 | 0.4306 | 0.998 | 11 | 239 | 83 | 10;3;3;11 |
| Seringhaus | ACT1 | staggered | 4 | 6;12;18;24 | -0.6444 | 0.6444 | 0.998 | 11 | 164 | 51 | 4;3;3;3;11 |
| Seringhaus | ACT1 | staggered | 5 | 6;12;18;24;30 | -0.8582 | 0.8582 | 0.998 | 5 | 77 | 13 | 4;3;3;3;3;5 |
| Seringhaus | HBG2 | centered | 3 | 17;18;19 | -0.1114 | 0.1114 | 0.974 | 16 | 481 | 196 | 15;16 |
| Seringhaus | HBG2 | centered | 5 | 16;17;18;19;20 | -0.24579999999999996 | 0.24579999999999996 | 0.974 | 15 | 421 | 169 | 14;15 |
| Seringhaus | HBG2 | centered | 7 | 15;16;17;18;19;20;21 | -0.3802 | 0.3802 | 0.974 | 14 | 365 | 144 | 13;14 |
| Seringhaus | HBG2 | centered | 9 | 14;15;16;17;18;19;20;21;22 | -0.5146 | 0.5146 | 0.974 | 13 | 313 | 121 | 12;13 |
| Seringhaus | HBG2 | centered | 11 | 13;14;15;16;17;18;19;20;21;22;23 | -0.649 | 0.649 | 0.974 | 12 | 265 | 100 | 11;12 |
| Seringhaus | HBG2 | staggered | 2 | 12;24 | -0.1673 | 0.1673 | 0.997 | 11 | 302 | 109 | 10;9;11 |
| Seringhaus | HBG2 | staggered | 3 | 12;18;24 | -0.39 | 0.39 | 0.997 | 11 | 239 | 83 | 10;3;3;11 |
| Seringhaus | HBG2 | staggered | 4 | 6;12;18;24 | -0.6127 | 0.6127 | 0.997 | 11 | 164 | 51 | 4;3;3;3;11 |
| Seringhaus | HBG2 | staggered | 5 | 6;12;18;24;30 | -0.8354000000000001 | 0.8354000000000001 | 0.997 | 5 | 77 | 13 | 4;3;3;3;3;5 |

### Table S4. Deng even-versus-random mismatch aggregate values

Reported aggregate relative signal values extracted from Deng et al. Figure 1/text; ratio calculated here.

| study | mismatches | evenly_distributed_relative_signal | randomly_distributed_relative_signal | random_to_even_signal_ratio |
|---|---|---|---|---|
| Deng | 0 | 1.0 | 1.0 | 1.0 |
| Deng | 1 | 0.7 | 0.7 | 1.0 |
| Deng | 2 | 0.36 | 0.52 | 1.4444444444444446 |
| Deng | 3 | 0.23 | 0.33 | 1.434782608695652 |
| Deng | 4 | 0.11 | 0.26 | 2.3636363636363638 |
| Deng | 5 | 0.04 | 0.12 | 3.0 |
| Deng | 6 | 0.02 | 0.11 | 5.5 |
| Deng | 7 | 0.01 | 0.09 | 9.0 |

### Table S5. Deng MPDNN model progression

Reported model correlations from Deng et al.; R2 calculated as R squared.

| model | R | note | R2 |
|---|---|---|---|
| Original scalar NN relative free energy | 0.798 | published Deng model baseline | 0.636804 |
| Positional weights added (topology) | 0.828 | thermodynamics with position weights | 0.685584 |
| Mismatch-dimer parameters adjusted | 0.921 | surface-calibrated topology-aware thermodynamics | 0.848241 |
| Matched + mismatch dimers adjusted | 0.938 | final MPDNN model | 0.8798439999999998 |

### Table S6. Rennie maximum perfect-match-length summary

Published summary values from Rennie et al. with no row-level reanalysis.

| feature | r | R2_or_variance_fraction | note |
|---|---|---|---|
| Maximum perfect-match length, 1-2 mismatch probes | 0.65 | 0.43 | Lmax explains 43% of variance in paired 1-MM vs 2-MM log2 ratios |
| Mismatch position variance contribution | | 0.046 | Position accounted for 4.6% of total variation |
| Substitution type variance contribution | | 0.0095 | Substitution type accounted for 0.95% of total variation |

### Table S7. Mechanistic and thermodynamic bridge evidence

Extracted summary evidence from Naiser et al. and Hooyberghs/Carlon; used for interpretation, not a pooled regression.

| Source | Mechanistic point | Numerical/detail reported | How used |
|---|---|---|---|
| Naiser et al. | Mismatch discrimination is mostly determined by defect position and sequence context | Central mismatches in 16-mer examples typically reduced signal to 0-40% of perfect match | Supports position/topology dependence |
| Naiser et al. | Double-ended zipper/partition-function model | Partition function over partially denatured states explains positional influence | Physical basis for ZboxNN |
| Naiser et al. | PDNN inferred from zipper model | Position-dependent nearest-neighbor weighting can be derived from partial denaturation | Connects topology to nearest-neighbor thermodynamics |
| Hooyberghs/Carlon | Microarray thermodynamics can correlate with solution thermodynamics | Pearson correlation 0.839 between microarray and solution DeltaDeltaG in their analysis | Shows scalar thermodynamics is useful but platform-dependent |

### Table S8. Affymetrix fixed-mismatch control rows

Row-level illustrative control retained from the ECI analysis; sequence-resolved source extraction should be deposited for final submission.

| Probe_ID | Mismatch_pattern | MM | Log2_intensity | DeltaG37_kcal_mol | Retained_cores | S_ECI | N_box_k2 | N_box_k3 | N_box_k4 | N_box_k5 | N_box_k6 |
|---|---|---|---|---|---|---|---|---|---|---|---|
| PM-A1 | Perfect match | 0 | 12.8 | -27.1 | 14 | 196 | 91 | 78 | 66 | 55 | 45 |
| CL-1 | Clustered | 1 | 12.4 | -25.4 | 20 | 400 | 190 | 171 | 153 | 136 | 120 |
| CL-2 | Clustered | 2 | 11.6 | -22.4 | 14+5 | 221 | 101 | 84 | 69 | 56 | 45 |
| CL-3 | Clustered | 3 | 10.9 | -18.7 | 12+4 | 160 | 72 | 58 | 46 | 36 | 28 |
| CL-4 | Clustered | 4 | 9.8 | -17.9 | 11 | 121 | 55 | 45 | 36 | 28 | 21 |
| DS-1 | Distributed | 2 | 8.7 | -22.3 | 5+4 | 41 | 16 | 9 | 4 | 1 | 0 |
| DS-2 | Distributed | 3 | 6.4 | -19.0 | 4+3 | 25 | 9 | 4 | 1 | 0 | 0 |
| DS-3 | Distributed | 3 | 5.8 | -18.8 | 3+3 | 18 | 6 | 2 | 0 | 0 | 0 |
| DS-4 | Distributed | 3 | 4.9 | -18.6 | 6 | 36 | 15 | 10 | 6 | 3 | 1 |
| DS-5 | Distributed | 3 | 4.4 | -18.7 | none | 0 | 0 | 0 | 0 | 0 | 0 |

### Table S9. Affymetrix topology and thermodynamic correlation summary

Pearson correlations and p-values computed from Table S8 rows.

| Subset | Predictor | n | Pearson_r | p_value |
|---|---|---|---|---|
| 2-4 mismatch subset | S_ECI | 8 | 0.9152814652470628 | 0.0014251613930117 |
| 2-4 mismatch subset | DeltaG37_kcal_mol | 8 | -0.4441113804158873 | 0.2703052580759403 |
| 2-4 mismatch subset | N_box_k3 | 8 | 0.8957690182009062 | 0.0026142485739281 |
| 2-4 mismatch subset | N_box_k4 | 8 | 0.8828879647420768 | 0.0036711054527036 |
| 2-4 mismatch subset | N_box_k5 | 8 | 0.8721008715692937 | 0.0047415970967068 |
| strict 3-mismatch subset | S_ECI | 5 | 0.9583418102580634 | 0.0101426608719099 |
| strict 3-mismatch subset | DeltaG37_kcal_mol | 5 | -0.0420103087197219 | 0.9465265513606884 |
| strict 3-mismatch subset | N_box_k3 | 5 | 0.943616978091154 | 0.0159348829759425 |
| strict 3-mismatch subset | N_box_k4 | 5 | 0.937794306861185 | 0.0184495040562741 |
| strict 3-mismatch subset | N_box_k5 | 5 | 0.9421569137484184 | 0.0165541392819907 |

### Table S10. HPV sequence-resolved edge-case audit

Sequence-resolved applied audit from the ambient HPV benchmark. Rows marked DISCREPANCY_CHECK should be verified before relying on them for probe ordering.

Table S10 is shown here as key numerical/audit columns. The full sequence-resolved table, including probe and off-target windows, is provided in the CSV and Excel workbook.

| case | role | s_eci_on | s_eci_off | delta_s_eci | exploratory_delta_delta_g37 | status | comment |
|---|---|---|---|---|---|---|---|
| CP1/HPV6 | clean concordant positive control | 104 | 16 | 88 | 8.164 | MATCHES_FIGURE | Shows that a non-intended sequence can retain local similarity but still be topologically fragmented relative to the intended retained island. |
| CP33/HPV70 | topology-strong, thermodynamic margin near uncertainty zone | 64 | 16 | 48 | -0.583 | MATCHES_FIGURE | Useful vignette: topology-defined strongest ECI off-target differs from the sequence-similarity/NN off-target, illustrating why the layers should be reported separately. |
| CP34/HPV72 | topology favorable; FASTA/Figure transcription discrepancy flagged | 64 | 25 | 39 | 2.068 | DISCREPANCY_CHECK | The sequence audit flags a difference between the FASTA-derived intended window and the manually transcribed Figure 5 target; keep as audit material until verified. |
| CP29/HPV66 | original thermodynamics-favorable/topology-neutral case; sequence-audit caution | 81 | 36 | 45 | 3.717 | MATCHES_FIGURE; LOW_CONFIDENCE_MANUAL_READ | The older manuscript used CP29 as topology-neutral; the sequence audit gives a different topology score. Keep CP29 as a discrepancy/audit case, not as a primary proof example. |
| CP9/HPV33 | weak topology margin with thermodynamically competitive non-intended sequence | 45 | 36 | 9 | -0.088 | MATCHES_FIGURE | A useful calibration case: topology is only weakly favorable and NN comparison is close/competitive. |
| CP5/HPV18 | boundary case; empirical exception; sequence discrepancy flagged | 16 | 9 | 7 | -2.700 | DISCREPANCY_CHECK | The original benchmark flagged CP5 as a review/calibration case; sequence audit also flags Figure/FASTA discrepancy and should not be used as primary validation. |
| CP21/HPV54 | strong boundary/failure mode; sequence discrepancy flagged | 13 | 68 | -55 | -5.369 | DISCREPANCY_CHECK | A boundary case: the off-target retains a longer topology island than the intended alignment. Use as a limitation/calibration example. |

Full columns in the machine-readable table: case, role, cp_seq, intended_segment, cp_vs_intended_pattern, eci_on_lengths, s_eci_on, strongest_eci_off_target, strongest_eci_off_window, cp_vs_strongest_eci_off_pattern, eci_off_lengths, s_eci_off, delta_s_eci, top_homology_target, top_homology_window, most_stable_nn_off_target, most_stable_nn_off_window, exploratory_delta_delta_g37, status, comment

### Table S11. Temperature/stringency calibration table

Calibration logic derived from Deng et al. temperature thresholds and the proposed room-temperature LFA use case.

| Condition | Published/derived observation | Implication for k_eff |
|---|---|---|
| 50 C, 50% formamide | Deng et al. reported that 3 evenly distributed mismatches reached the practical low-signal criterion | Higher stringency supports larger effective threshold |
| 45 C, 50% formamide | Deng et al. reported that 4 evenly distributed mismatches were needed | Intermediate stringency |
| 42 C, 50% formamide | Deng et al. reported that 5 evenly distributed mismatches were needed | Lower stringency requires more fragmentation |
| Room temperature post-PCR LFA | Not directly measured in this manuscript | Must be calibrated empirically in final strip/buffer because shorter boxes may survive |

#### Supplementary References


1. SantaLucia,J. Jr and Hicks,D. (2004) The thermodynamics of DNA structural motifs. Annu. Rev. Biophys. Biomol. Struct., 33, 415-440.

2. Seringhaus,M., Rozowsky,J., Royce,T., Nagalakshmi,U., Jee,J., Snyder,M. and Gerstein,M. (2008) Mismatch oligonucleotides in human and yeast: guidelines for probe design on tiling microarrays. BMC Genomics, 9, 635.

3. Deng,Y., He,Z., Van Nostrand,J.D. and Zhou,J. (2008) Design and analysis of mismatch probes for long oligonucleotide microarrays. BMC Genomics, 9, 491.

4. Rennie,C., Noyes,H.A., Kemp,S.J., Hulme,H., Brass,A. and Hoyle,D. (2008) Strong position-dependent effects of sequence mismatches on signal ratios measured using long oligonucleotide microarrays. BMC Genomics, 9, 317.

5. Naiser,T., Kayser,J., Mai,T., Michel,W. and Ott,A. (2008) Position dependent mismatch discrimination on DNA microarrays - experiments and model. BMC Bioinformatics, 9, 509.

6. Hooyberghs,J., Van Hummelen,P. and Carlon,E. (2010) The effects of mismatches on hybridization in DNA microarrays: determination of nearest neighbor parameters. arXiv:1001.0653.

7. Hadiwikarta,W.W., Walter,J.C., Hooyberghs,J. and Carlon,E. (2012) Probing hybridization parameters from microarray experiments: nearest-neighbor model and beyond. Nucleic Acids Res., 40, e138.

8. Harbig,J., Sprinkle,R. and Enkemann,S.A. (2005) A sequence-based identification of the genes detected by probesets on the Affymetrix U133 plus 2.0 array. Nucleic Acids Res., 33, e31.

9. Wang,Y., Miao,Z.H., Pommier,Y., Kawasaki,E.S. and Player,A. (2007) Characterization of mismatch and high-signal intensity probes associated with Affymetrix GeneChips. Bioinformatics, 23, 2088-2095.

10. Brukner,I., El-Ramahi,R., Sawicki,J., Gorska-Flipot,I., Krajinovic,M. and Labuda,D. (2007) Hybridization assay performed at ambient temperature for typing high-risk human papillomaviruses. J. Clin. Virol., 39, 113-118.

11. Brukner,I., El-Ramahi,R., Gorska-Flipot,I., Krajinovic,M. and Labuda,D. (2007) An in vitro selection scheme for oligonucleotide probes to discriminate between closely related DNA sequences. Nucleic Acids Res., 35, e66.

12. Brukner,I., Krajinovic,M., Dascal,A. and Labuda,D. (2007) A protocol for the in vitro selection of specific oligonucleotide probes for high-resolution DNA typing. Nat. Protoc., 2, 2806-2819.

13. Baker,G.C., Smith,J.J. and Cowan,D.A. (2003) Review and re-analysis of domain-specific 16S primers. J. Microbiol. Methods, 55, 541-555.

14. Klindworth,A., Pruesse,E., Schweer,T., Peplies,J., Quast,C., Horn,M. and Glockner,F.O. (2013) Evaluation of general 16S ribosomal RNA gene PCR primers for classical and next-generation sequencing-based diversity studies. Nucleic Acids Res., 41, e1.
